# Comparative mitogenomics of native European and alien Ponto-Caspian amphipods

**DOI:** 10.1101/2023.03.15.532755

**Authors:** Jan-Niklas Macher, Eglė Šidagytė-Copilas, Denis Copilaş-Ciocianu

## Abstract

European inland surface waters harbor an extensive diversity of native amphipod crustaceans with many species facing threats from invasive counterparts of Ponto-Caspian origin. Herein, we examine mitochondrial genomes to infer phylogenetic relationships and compare gene order and nucleotide composition between representative native European and invasive Ponto-Caspian taxa belonging to five families, ten genera, and 20 species (13 newly sequenced herein). We observe diverse gene rearrangement patterns in the phylogenetically disparate native species pool. *Pallaseopsis quadrispinosa* and *Synurella ambulans* show significant departures from the typical organization, with extensive translocations of tRNAs and the nad1 gene, as well as a tRNA-F polarity switch in the latter. The monophyletic alien Ponto-Caspian gammarids display a conserved gene order, mainly differing from the native species by a tRNA-E and tRNA-R translocation, which strengthens previous findings. However, extensive rearrangement is observed in *Chaetogammarus warpachowskyi* with translocations of six tRNAs. The alien corophiid, *Chelicorophium curvispinum*, displays a very conserved gene order despite its distant phylogenetic position. We also find that native species have a significantly higher GC and lower AT content than invasive ones. The observed mitogenomic differences between native and invasive amphipods need further investigation and could shed light on the mechanisms underlying invasion success.

## 1. Introduction

The European continent harbors a great diversity of inland amphipod crustaceans, both in surface or subterranean, fresh or brackish waters [1–4]. Moreover, this diversity is significantly underestimated due to the extensive prevalence of cryptic species [5–8]. Nevertheless, an important proportion of this fauna is under threat due to invasive species, climate change, eutrophication and other anthropogenic factors [9–12]. One of the main threats that native surface amphipods face is competition and eventual extinction due to the spread of invasive counterparts, especially originating from the Ponto-Caspian basin. [9,13–15].

The Ponto-Caspian region encompasses the Azov, Black, Caspian and Aral seas as well as the lower stretches of their tributaries [13]. The area is characterized by a peculiar endemic fauna, particularly adapted to broad salinity fluctuations [16,17]. Due to this environmental tolerance, many Ponto-Caspian endemics became invasive by expanding their range beyond the native borders, mainly colonizing European inland waters, but also making their way to other continents [13,18]. One of the most successful groups are amphipods, with up to 40% of the species pool spreading outside the native range during the last century mainly due to increases in shipping activity, construction of canals, and intentional introductions [13,19]. The invasive Ponto-Caspian amphipods are competitively superior to the native species they encounter along invasion routes, leading to their decline and eventual extinction [20–25].

Comparative studies involving both native and invasive species are necessary in order to understand invasion success. However, the underlying molecular and genetic mechanisms behind the success of Ponto-Caspian species are poorly known and studies are in their infancy [26,27]. The mitochondrial genome is a good candidate for such comparative molecular studies as mitochondria are fundamental for the functioning of multicellular life and complete mitochondrial genomes are relatively cheap and easy to sequence due to recent advances in high-throughput sequencing and bioinformatic pipelines [28,29]. Given that studies generally reveal strong differentiation in respiratory function among native and invasive aquatic species [30–33], it is sensible to presume that the structure of the mitochondrial genome may hold insight regarding invasion success.

To date, relatively few mitochondrial genomes are available for the invasive Ponto-Caspian amphipods and for native European species, many of which were obtained from transcriptomic data and are thus of varying reliability [27,34–37]. Of the 13 widespread invasive Ponto-Caspian species, the mitochondrial genomic structure is reliably known in four species from two genera (*Dikerogammarus bispinosus, D. haemobaphes, D. villosus*, and *Pontogammarus robusotides*) while the mitogenome of *Obesogammarus crassus* is only known from transcriptomic reads, thus some regions have low coverage and reliability [27]. With respect to the native European species, the situation is more severe because only four species out of several dozens (possibly hundreds) have reliably known mitogenomes (*Gammarus duebeni, G. fossarum, G. lacustris*, and *G. roeselii*)[34–36,38] while four more species have mitogenomes assembled from transcriptomic reads (*G. pulex, G. wautieri, Echinogammarus berilloni* and *Pectenogammarus veneris*)[27,39].

In this study we compare the mitochondrial gene order, nucleotide composition and assess the phylogenetic relationships of native European and invasive Ponto-Caspian amphipods. We present a significantly expanded dataset that includes mitochondrial genomes representing most major native and invasive species in Europe. We present the first mitochondrial genomes of the native *Synurella ambulans, Pallaseopsis quadrispinosa, G. jazdzewskii*, and *G. varsoviensis*, the first DNA-based mitogenome for *G. pulex*, and the first mitogenome of *G. lacustris* from Europe (previously sequenced only from the Tibetan Plateau [38]). Regarding the invasive species, we present the first mitogenomes for *Chaetogammarus ischnus* and *C. warpachowskyi*, the first DNA-based mitogenome for *O. crassus*, and additional mitogenomes for *D. haemobaphes, D. villosus* and *P. robustoides*. Lastly, we present the first mitogenome for *Chelicorophium curvispinum*, the most widespread Ponto-Caspian corophiid amphipod.

## 2. Material and Methods

### 2.1. Sampling, laboratory protocols and sequencing

Animals used in the analyses were collected from Lithuania, Poland, and Latvia between 2018 and 2020 using kick-sampling with a hand net (see Table S1 for detailed locality information). Specimens were stored in 96% ethanol in the field. Afterward, the ethanol was exchanged several times, and the material was stored at -20°C. Specimens were identified using relevant keys [4,40,41]. Th taxonomy of the focal taxa follows the most recent updates [4,42–45].

We dissected the dorsal half of the animal (from head to urosome) using microsurgical scissors and fine needles to avoid contamination from the gut, and extracted genomic DNA using the Quick-DNA Miniprep Plus Kit (Zymo Research) with the lysis step prolonged overnight. All specimens selected for high-throughput sequencing were also DNA barcoded using the protocols described in [46] to further confirm morphological identifications.

After DNA extraction, we assessed quantity and fragment length of the genomic DNA using a FragmentAnalyzer (Agilent, USA). To fragment the DNA, the Covaris M220 system (Covaris, UK) was used targeting a fragment size of 250 base pairs. The fragmented DNA was then checked again on the FragmentAnalyzer system to confirm the quantity and length of fragments. The NEBNext Ultra II DNA Library Prep Kit and corresponding NEBNext Multiplex Oligos for Illumina were used to prepare shotgun genomic libraries following the manufacturer’s protocol. The final library concentration and fragment size were confirmed on a TapeStation (Agilent) before manual equimolar pooling of samples. A negative control was processed together with the samples and did not show any DNA. The final library was sequenced using the Illumina NovaSeq 6000 platform with 2×150 bp read length at Macrogen Europe.

### 2.2. Bioinformatics, mitochondrial genome assembly and annotation

Raw data were checked for low-quality samples using the FastQC software and Illumina adapters were trimmed using Trimmomatic [47]. Strict quality filtering was applied to trimmed reads using vsearch, with reads truncated at the first base with a phred score <15. Reads shorter than 100bp were excluded from subsequent analysis. Per sample, ten million quality-checked reads were assembled using Megahit [48] on the Naturalis high-performance cluster, with kmer lengths ranging from 15 to 115. The resulting contigs were imported into Geneious Prime (v.2022.1), and BLAST searches were conducted against a manually compiled reference library of amphipod mitochondrial genes [27,36] downloaded from NCBI GenBank (https://www.ncbi.nlm.nih.gov/genbank/). Contigs were identified as potential mitochondria based on BLAST results and contig lengths (between 10,000 and 20,000bp). Potential mitochondrial contigs were subsequently annotated using Mitos2 [49]. Annotations were manually checked and refined in Geneious Prime, and gene sequences (nucleotide and protein) were extracted for subsequent phylogenetic analyses.

### 2.3. Nucleotide composition

To the 13 mitogenomes obtained in this study we added 7 mitogenomes obtained in previous studies, totaling 20 species, of which 8 were Ponto-Caspian invaders and 12 native species (Table1). Nucleotide composition was calculated for the entire mitogenomes using MEGA 6 [50]. In order to visualize patterns of composition among species, the percentage matrix of each of the four nucleotides was subjected to a Principal Component Analysis (PCA) using a variance-covariance matrix. A Permutational Multivariate Analysis of Variance (PERMANOVA) test with 9999 permutations was used to detect differences in nucleotide composition among the native and invasive species groups. Furthermore, GC and AT content were separately compared among native and invasive species using a Mann-Whitney test. All analyses were conducted with PAST 4.10 [51].

**Table 1.** Overview of the species used in the comparative analyses.

| Family | Species | NCBI accession number | Status | Mitogenome length (bp) | A % | T % | G % | C % | Source |
| --- | --- | --- | --- | --- | --- | --- | --- | --- | --- |
| Corophiidae | *Chelicorophium curvispinum*<sup>1</sup> | CC6 | Invasive | 14867 | 37.8 | 30.6 | 12.6 | 19.1 | This study |
| Gammaridae | *Chaetogammarus warpachowskyi*<sup>1</sup> | CW4 | Invasive | 17336 | 35.2 | 35.9 | 10.9 | 18.0 | This study |
| Gammaridae | *Chaetogammarus ischnus*<sup>1</sup> | EI4 | Invasive | 14694 | 32.5 | 33.9 | 12.1 | 21.5 | This study |
| Gammaridae | *Dikerogammarus bispinosus* | OK173840 | Invasive | 15336 | 33.9 | 36.6 | 11.1 | 18.3 | [27] |
| Gammaridae | *Dikerogammarus haemobaphes*<sup>1</sup> | DH3 | Invasive | 15258 | 31.9 | 34.2 | 13.1 | 20.9 | This study* |
| Gammaridae | *Dikerogammarus villosus*<sup>1</sup> | DV4 | Invasive | 15176 | 32.7 | 35.1 | 12.3 | 19.9 | This study* |
| Pontogammaridae | *Obesogammarus crassus*<sup>1</sup> | OC4 | Invasive | 15838 | 33.6 | 37.5 | 11.3 | 17.6 | This study† |
| Pontogammaridae | *Pontogammarus robustoides*<sup>1</sup> | PR4 | Invasive | 15917 | 33.3 | 36.2 | 11.8 | 18.7 | This study* |
| Crangonyctidae | *Synurella ambulans* | SA1 | Native | 15652 | 32.2 | 30.8 | 13.3 | 23.6 | This study |
| Gammaridae | *Gammarus duebeni* | JN704067 | Native | 15651 | 32.5 | 22.0 | 31.5 | 14.0 | [35] |
| Gammaridae | *Gammarus fossarum* | KY197961 | Native | 15989 | 32.0 | 22.0 | 33.2 | 12.9 | [36] |
| Gammaridae | *Gammarus lacustris* | GL1 | Native | 18195 | 31.1 | 32.8 | 13.3 | 22.8 | This study* |
| Gammaridae | *Gammarus jazdzewskii* | GZ1 | Native | 16136 | 34.6 | 34.4 | 11.4 | 19.5 | This study |
| Gammaridae | *Gammarus pulex* | GP2 | Native | 14886 | 33.1 | 34.0 | 12.2 | 20.7 | This study† |
| Gammaridae | *Gammarus roeselii* | MG779536 | Native | 16073 | 33.9 | 32.9 | 12.3 | 20.9 | [34] |
| Gammaridae | *Gammarus varsoviensis* | GV1 | Native | 15482 | 31.1 | 32.8 | 13.2 | 22.8 | This study |
| Gammaridae | *Gammarus wautieri* | BK059229 | Native | 13927 | 32.4 | 22.2 | 34.2 | 11.2 | [27,39]† |
| Gammaridae | *Echinogammarus berilloni* | BK059223 | Native | 14454 | 30.2 | 26.9 | 28.0 | 14.9 | [27,39]† |
| Gammaridae | *Pectenogammarus veneris* | BK059233 | Native | 14369 | 34.1 | 22.2 | 31.4 | 12.4 | [27,39]† |
| Pallaseidae | *Pallaseopsis quadrispinosa* | PQ1 | Native | 16147 | 30.9 | 30.9 | 15.3 | 22.9 | This study |

<sup>1</sup> – Ponto-Caspian species; \* – species whose mitogenomes were also sequenced in previous studies; † – species whose mitogenomes were previously assembled from RNA sequences.

### 2.4. Phylogenetic analyses

The purpose of these analyses was to place the focal taxa within the wider phylogenetic context of the Amphipoda. Altogether, the data obtained in this study was merged with additional 62 mitogenomes from the literature representing 25 families and 59 species, including one isopod outgroup, *Ligia oceanica* (see Supplementary Table S1 for further details). The analyses were based on the 13 protein coding genes and excluded the large (16S rRNA) and small (12S rRNA) ribosomal subunits. Protein-coding genes evolve in a more predictable fashion then the erratic ribosomal units and can confidently be aligned. Each of the 13 genes were aligned separately by codon using MUSCLE [52] implemented in MEGA 6 with default options. All nucleotide alignments were protein translated using the invertebrate mitochondrial genetic code (translation table 5). Individual gene alignments were concatenated using SequenceMatrix [53]. Both nucleotide and translated protein sequences were used in the phylogenetic analyses. The concatenated nucleotide matrix had a total length of 11047 bp, while the protein matrix was 3682 amino acids long. Best partitioning schemes (by codon) and evolutionary models for the nucleotide data were selected with PartitionFinder 2 [54].

Phylogenetic analyses were conducted within a Bayesian (BI) framework with Phylobayes MPI 1.8c [55], and a maximum-likelihood (ML) framework with IQ-Tree 2.1.2 [56]. Phylobayes nucleotide analyses were run for 10000 cycles using the GTR exchange rates and the CAT profile mixture. Convergence, mixing, and effective samples size were checked by examining the relative difference among chains (<0.2) as well as using Tracer 1.7 [57]. IQ-Tree nucleotide analyses were run under an edge-linked model with each partition having an independent evolutionary model selected with PartitionFinder 2. Node support was assessed using 1000 ultrafast bootstrap replicates [58]. The protein phylogenetic analyses were run with the general metazoan mitochondrial amino acid substitution model (MtZOA)[59] for both Phylobayes and IQ-tree with the same settings as for nucleotides. All phylogenetic analyses were carried out using the computational infrastructure available at the CIPRES Science Gateway [60].

## 3. Results

### 3.1. Mitochondrial genomic structure

All samples resulted in high-quality reads that could be assembled into full mitochondrial genomes containing the expected number of 13 protein-coding genes, large and small-subunit rRNA and 22 tRNAs. In most cases the inferred gene order is similar between the native and invasive Ponto-Caspian species. The most commonly observed difference is a translocation of the tRNA-E and tRNA-R, which is in line with previous observations based on less extensive taxonomic datasets [27,37]. However, there are also a few rather contrasting patterns of variation between the native and invasive groups (Fig. 1). The native species exhibit three general patterns: (1) minor translocations (swaps) between transfer RNAs (tRNA-N, tRNA-E, and tRNA-R) as observed in *G. roeselii* and *G. varsoviensis*; (2) significant translocations of several tRNAs and the NADH dehydrogenase 1 gene (nad1) in *P. quadrispinosa* and *S. ambulans*; (3) a polarity switch of the tRNA-F in *S. ambulans*. The gene arrangement in the Ponto-Caspian gammarids is identical in all studied species except *C. warpachowskyi*, which shows a significant departure with translocation of six tRNAs. The gene order in the Ponto-Caspian alien corophiid *C. curvispinum* is identical to that of most native species (Fig. 1). In general, the gene arrangements seem to follow phylogenetic relationships.

**Fig. 1.**
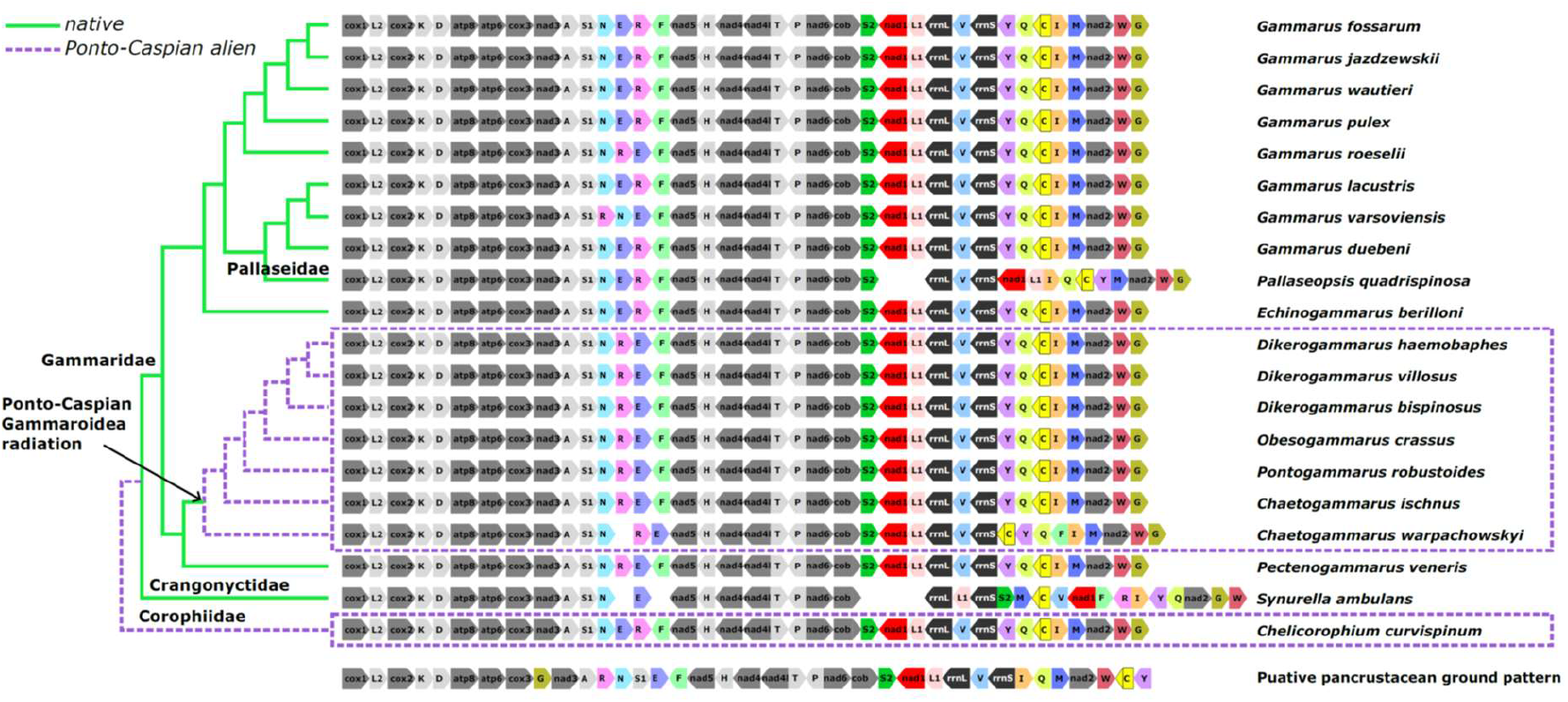
Comparison of mitochondrial genome organization among focal species. Genes with conserved positions are shown in grayscale, translocated genes are colored. Genes encoded on the plus-strand point towards the right, while the ones encoded on the minus-strand point towards the left. Invasive Ponto-Caspian species are shown with dashed lines. The cladogram on the left represents a summary of phylogenetic relationships form Fig. 3.

### 3.2. Nucleotide composition

Multivariate analyses indicate a significant differentiation with respect to nucleotide composition between the native and invasive species. The PCA scatterplot indicates a modest overlap between native and invasive groups in multivariate space, with the first two axes explaining 98% of the observed variance (Fig. 2A). The invasive species are generally associated with a higher AT content while native species with a higher GC content, although a large variation in GC content of native species is observed. The separation is further confirmed by PERMANOVA testing which indicates significant differences among the two groups (F = 6.257, *p* = 0.01). Further univariate comparisons using Mann-Whitney tests reveal that invasive species have a significantly higher AT and significantly lower GC content that the native species (mean AT% _invasive_ = 68.86, mean AT% _native_ = 60.99, z = 3.04, *p* = 0.001; mean GC% _invasive_ = 31.15, mean GC% _native_ = 38.97, z = 3.48, *p* = 0.001) (Fig. 2B).

**Fig. 2.**
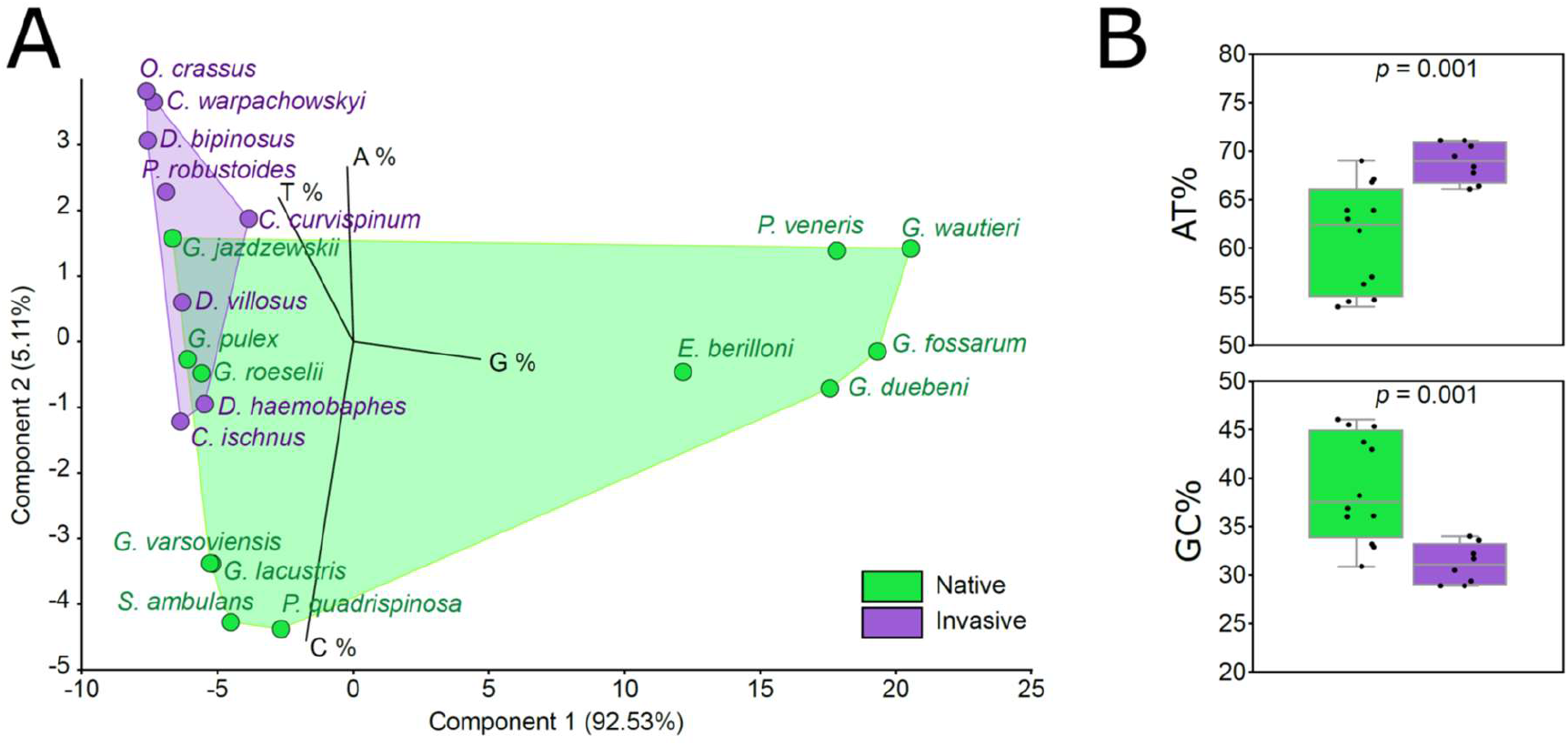
Differentiation of native European and invasive Ponto-Caspian species with respect to nucleotide composition across the entire mitochondrial genome. A) PCA scatterplot depicting multivariate differentiation across all four nucleotides, B) boxplots comparing AT and GC content among native and invasive species.

### 3.3. Phylogenetic analyses

Phylogenetic analyses revealed congruent relationships among methods (BI and ML) and datasets (nucleotides and amino acids) (Fig. 3). Disagreements were observed only at unsupported nodes. The native European inland water species are phylogenetically diverse, being interspersed among two main superfamilies, the Gammaroidea and Crangonyctoidea. Although the alien Ponto-Caspian species also belong to two main superfamilies, Gammaroidea and Corophioidea, the gammarids form a strongly supported monophylum. Our analyses reveal for the first time the phylogenetic position of *P. quadrispinosa*, confirming it as a sister species to the Baikal endemic *P. kessleri* and ultimately part of the Baikal Lake acanthogammarid radiation.

**Fig. 3.**
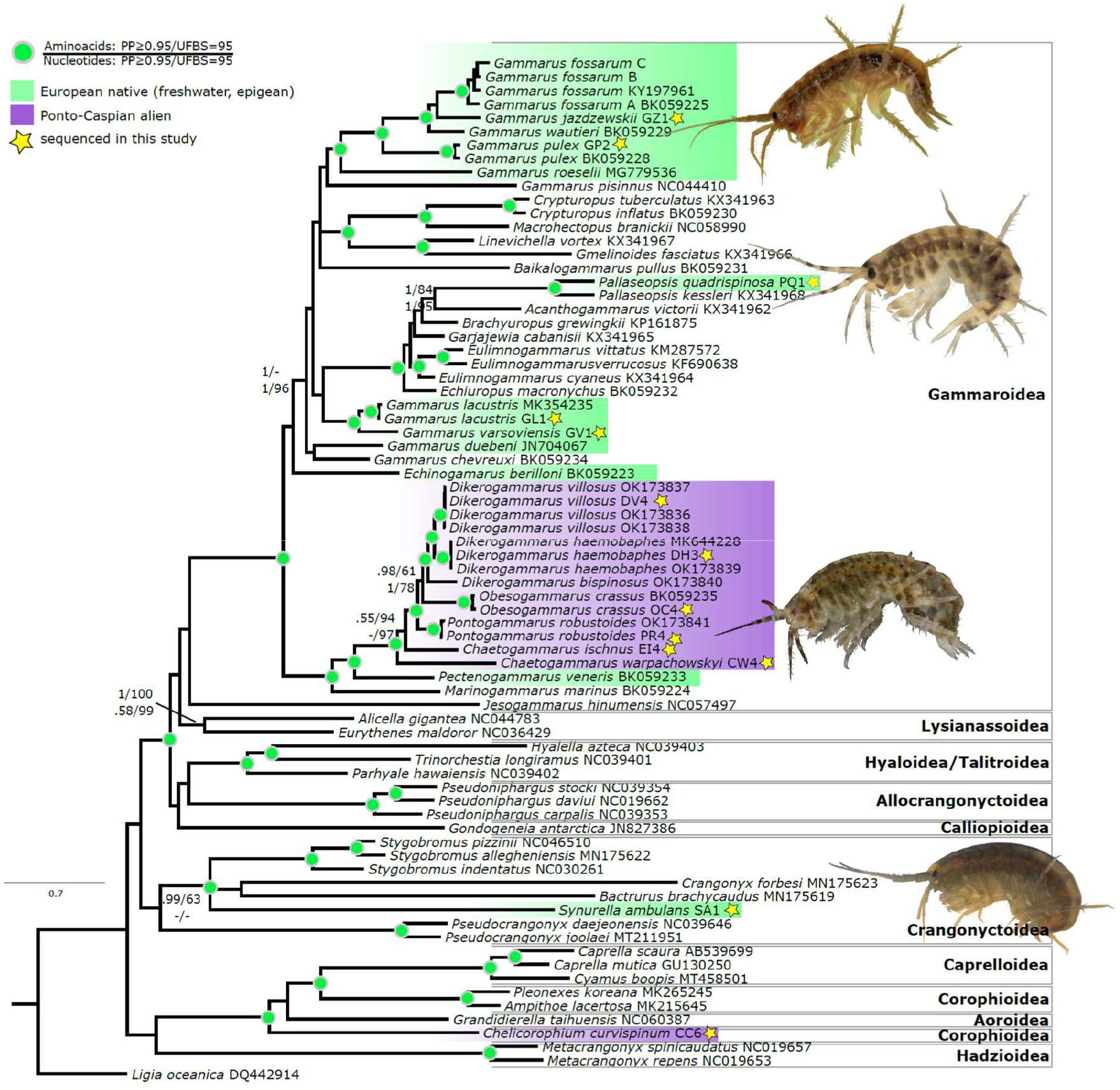
Amino acid Bayesian phylogeny based 13 mitochondrial protein coding genes depicting the evolutionary relationships among the focal taxa (highlighted with color). Native European surface-dwelling species are shown with green shading, while invasive Ponto-Caspian species are in purple. Stars indicate taxa sequenced in this study. Green circles indicate nodes that received strong support in all analyses. Nodes with numbers received moderate to strong support. Numbers above nodes indicate statistical support (posterior probabilities—PP; ultrafast bootstrap—UFBS) for amino acid-based trees; below nodes for nucleotide-based trees. Nodes that are not annotated received weak/no support. Inset photographs from top to bottom: *G. fossarum, P. quadrispinosa, C. warpachowskyi*, and *S. ambulans* (D. Copilaş-Ciocianu).

## 4. Discussion

The patterns of mitogenomic rearrangements observed in this study are consistent with the diversity that has been observed in other amphipod clades, ranging from major differentiation at generic levels to highly conserved between divergent clades [61–64]. The mitogenomic phylogenetic relationships obtained herein are also in agreement with other phylogenetic studies based on nuclear and mitochondrial markers [65].

Our study reveals that the native inland European amphipods can exhibit substantial departures from the typical mitogenomic organization, while the alien Ponto-Caspian species are more conservative. This is not unexpected given the greater phylogenetic disparity among the native species. However, the organization patterns seem not always phylogeny-driven. For example, *C. curvispinum*, which is distantly related to the other focal species in this study, exhibits a conserved gene arrangement, identical to that of most native species. On the other hand, *P. quadrispinosa* is more closely related to other native gammarids, yet it diverges significantly with respect to gene order. In fact, the gene order of *P. quadrispinosa* is identical to that of its congener from Lake Baikal, *P. kessleri* [66]. Our study confirms for the first time with molecular data the phylogenetic position of this species, which is a glacial relict that has almost gone extinct in Central Europe due to climate warming and eutrophication [11,67]. The peculiar mitogenomic structure of *Pallaseopsis* is outstanding even among other Baikalian amphipods [66,68–70], possibly reflecting intense periods of selection [71,72].

The native crangonyctid *S. ambulans* is phylogenetically very distant from the native gammarids and its mitogenomic structure is highly distinct as well, with several tRNAs and the nad1 gene having undergone translocations. Moreover, we detected a switch to a positive polarity of the tRNA-F gene, which normally is found on the minus-strand in amphipods. This pattern is partially phylogeny-driven, because the available mitogenomes of other crangonyctids seem to be generally conserved, but in some cases can show significant transpositions [73]. The remaining native gammarids (*Echinogammarus, Gammarus*, and *Pectenogammarus*) have a conserved mitogenomic structure, the main differences consisting in minor translocations of tRNAs, especially between tRNA-E and tRNA-R. The mitogenome of *G. varsoviensis* exhibits a previously unknown translocation of the tRNA-N, which is located between the tRNA-R and the tRNA-E.

The alien Ponto-Caspian gammarids have a much more conserved gene order than their native counterparts. All species, except the phylogenetically distant *C. warpachowskyi*, have identically structured mitogenomes. They differ from the native species by a swap between the tRNA-E and tRNA-R. This pattern has been previously observed on less taxonomically comprehensive datasets [27,37]. However, here we show that this does not apply to all Ponto-Caspian gammarids, as *C. warpachowskyi* shows significant departures from this pattern with translocations of six tRNAs. This could be due to phylogeny as this species is more distantly related to the other Ponto-Caspian gammarids and should be assigned to a new genus [46,74]. Sequencing of additional mitogenomes from Ponto-Caspian gammaroidean species will likely reveal even more patterns of gene rearrangements as only seven out of 82 species have been sequenced so far [4].

Apart from gene order, we found extensive differentiation with respect to nucleotide composition between the native and invasive Ponto-Caspian species. The invasives have significantly more AT rich mitogenomes than the natives, while natives have a higher GC content. This indicates that invasive species might have longer non-coding regions or that the native species have longer protein-coding genes, overall indicating more compact mitogenomes in the latter [75,76]. How and if this relates to invasion ability remains to be seen, but this strong differentiation can potentially open new avenues for research.

## 5. Conclusion

Our comparative analyses revealed a substantial differentiation between the mitogenomes of native European and invasive Ponto-Caspian amphipod crustacean species. The natives are more phylogenetically disparate, exhibit diverse mitogenomic configurations, and higher GC content than the invasives, which exhibit phylogenetic compactness, with highly conserved gene order and higher AT content. We suggest that these findings are not solely phylogeny-driven, as gene order conservation can vary across phylogenetic depths. Investigating the biological implications of the mitogenomic differences between native and invasive amphipods could shed light on the adaptive mechanisms underlying invasion success.

## Acknowledgements

This study was financed by the Research Council of Lithuania (contract no. S-MIP-20-26).

## Data accessibility

All raw reads are available in the NCBI Sequence Read Archive (SRA), accession numbers: (available upon acceptance). All mitochondrial genomes and annotations are available in NCBI GenBank, (accession numbers available upon acceptance) as well as on Figshare (DOI available upon acceptance).

